# Electrophysiological classification of *CACNA1G* gene variants associated with neurodevelopmental and neurological disorders

**DOI:** 10.1101/2025.03.18.644009

**Authors:** Amaël Davakan, Leos Cmarko, Barbara Ribeiro Oliveira-Mendes, Claire Bernat, Najlae Boulali, Jérôme Montnach, Stephanie E. Vallee, Lydie Burglen, Vincent Cantagrel, Norbert Weiss, Sophie Nicole, Arnaud Monteil, Michel De Waard, Philippe Lory

**Affiliations:** IGF, Université de Montpellier, CNRS, INSERM, Montpellier, France; LabEx ‘Ion Channel Science and Therapeutics’, France; Nantes Université, CNRS, INSERM, l’Institut du Thorax, Nantes, France; Department of Pathophysiology, Third Faculty of Medicine, Charles University, Prague, Czech Republic; Dartmouth-Hitchcock Medical Center and Clinics, Department of Pediatrics and Clinical Genetics, Lebanon, NH, USA; Université Paris Cité, INSERM UMR1163, Imagine Institute, Developmental Brain Disorders Laboratory, 75015, Paris, France; Pediatric Neurogenetics Laboratory, Department of Genetics, Armand-Trousseau Hospital, APHP, Sorbonne University, Paris, France

## Abstract

This study highlights the complementarity of automated patch-clamp (APC) and manual patch-clamp (MPC) approaches to describe the electrophysiological properties of eighteen Ca_v_3.1 calcium channel variants associated with various neurological conditions. Current density was measured efficiently for all variants in APC experiments, with four variants (p.V184G, p.N1200S, p.S1263A and p.D2242N) showing high current densities, compared to wild-type Ca_v_3.1 channel, while six variants (p.M197R, p.V392M, p.F956del, p.I962N, p.I1412T, and p.G1534D) displayed low current densities, and were therefore preferentially studied using MPC. The electrophysiological properties were well conserved in APC (e.g. inactivation and deactivation kinetics, steady-state properties), with only the APC-MPC correlation for the activation kinetics being less robust. In addition, neuronal modeling, using a deep cerebellar neuron (DCN) environment, revealed that most of the variants localized in the intracellular gate (S5 and S6 segments) could increase DCN spike frequencies. This DCN firing was critically dependent on the current density and further pointed to the gain-of-function (GOF) properties of p.A961T and p.M1531V, the recurrent variants associated with Spinocerebellar Ataxia type-42 with Neurodevelopmental Deficit (SCA42ND). Action-potential (AP) clamp experiments performed using cerebellar and thalamic neuron activities further established the GOF properties of p.A961T and p.M1531V variants. Overall, this study demonstrates that APC is well-suited to high-throughput analysis of Ca_v_3.1 channel variants, and that MPC complements APC for characterizing low-expression variants. Furthermore, *in silico* modeling and AP clamp experiments establish that the gain- or loss-of-function properties of the variants are determined by how the Ca_v_3.1 channel decodes the electrophysiological context of a neuron.

## Introduction

The T-type calcium channels (Ca_v_3.1, Ca_v_3.2 and Ca_v_3.3) are voltage-gated calcium channels with some unique features, being activated by low plasma membrane depolarization (low-voltage activated) and exhibiting fast inactivation (transient) (Zamponi et al., 2015;Lory et al., 2020). The Ca_v_3.1 channel is expressed in many types of neurons, from the cerebellum, thalamus and cortex, and contributes to neuronal excitability, especially rebound burst firing (Perez-Reyes, 2003). Several variants of *CACNA1G*, the gene encoding Ca_v_3.1 channels, are associated with neurological conditions, mainly cerebellar and neurodevelopmental, including SCA42 (spinocerebellar ataxia type 42; (Coutelier et al., 2015)), SCA42ND (SCA42 with neurodevelopmental deficits; (Chemin et al., 2018)), and developmental and epileptic encephalopathy (DEE; (Berecki et al., 2020)). Besides SCA42, most of these recently reported Ca_v_3.1 variants are missense *de novo* variants that strikingly alter the biophysical properties of the Ca_v_3.1 channel (Lory et al., 2020). This is well exemplified for the two first variants identified in SCA42ND patients, p.A961T and p.M1531V that localized in the intracellular gate (IG) in the IIS6 and IIIS6 segments, respectively, and are responsible for a slowing in the inactivation and deactivation kinetics and an increase of the window current (Chemin et al., 2018). Recently, several novel Ca_v_3.1 variants (p.M197R, p.V392M, p.F956del, p.I962N, p.S1263A, p.I1412T, p.G1534D and p.R1718G) were reported in patients with neurodevelopmental conditions, either fully or partly matching the original SCA42ND phenotype (Qebibo et al., 2024). All these findings have established *CACNA1G*, the gene encoding Ca_v_3.1 channels, as involved in a variety of neurological and neurodevelopmental diseases.

Patch-clamp recordings of Ca_v_3.1 variants heterologously expressed in HEK-293 cells is a gold standard for the electrophysiological characterization of disease variants (Chemin et al., 2018). However, facing the growing number of identified Ca_v_3.1 variants, calcium current recordings using manual patch-clamp (MPC) appear fastidious and there is a need for the use of medium/high-throughput electrophysiology for such characterization. Automated patch-clamp (APC) was recently employed for the investigation of sodium and potassium channel disease variants (Yajuan et al., 2012;Jiang et al., 2022;Vanoye et al., 2022;Ma et al., 2024). This experimental strategy was also used for the investigation of multiple variants in the Ca_v_3.3 channel associated with schizophrenia risk (Baez-Nieto et al., 2022). Not only can most MPC protocols be transposed to APC for accurate electrophysiological description of the recombinant Ca_v_3 channels, APC also offers specific advantages, such as parallel recordings of large number of cells (Montnach et al., 2021).

In this study, we provide the electrophysiological characterization of 18 Ca_v_3.1 variants, including 6 newly reported variants. APC and MPC approaches were performed jointly to validate the experimental conditions in APC experiments, i.e. the efficiency of transient transfection to record Ca_v_3.1 current, and the required adjustment of the external (5 mM *versus* 2 mM CaCl_2_) and intracellular (CsF versus CsCl) recording solutions. APC enabled accurate assessment of Ca_v_3.1 current density, while MPC was better suited to studying low-expressing variants. Neuronal modeling, using the deep cerebellar neuron (DCN) as well as action-potential clamp experiments were further carried out to functionally classify as loss-of-function (LOF) / unchanged / gain-of-function (GOF) this set of variants, pointing further to the clear GOF effect of p.A961T and p.M1531V, the two recurrent variants in SCA42ND.

## Materials and Methods

### Directed mutagenesis

The human *CACNA1G* complementary DNA (accession number NM_198387.2) was mutated to generate the 18 variants by using a site-directed mutagenesis service (GenScript Biotech, Netherlands). The protein variant nomenclature used here (e.g. p.A961T) is based on the UniProt protein sequence O43497 that corresponds to the full-length reference transcript, and is simplified in the Figures (as A961T) for an easier reading of the panels. The plasmid expression vectors (pcDNA3-based) were then amplified to reach the necessary dilution, 6 µg/µL for transfections for the APC experiments and 1 µg/µL for transfections for the MPC experiments.

### Transient transfection

Transient transfection was performed in 35 mm Petri dishes. For MPC, HEK-293T cells were transfected using jet-PEI (QBiogen) with a 2 µg plasmid DNA mix containing 1% of a GFP-encoding construct and 99% of a Ca_v_3.1-encoding construct, either wt or mutant channels. For APC, HEK-293 cells were transfected by electroporation using the MaxCyte STx system (MaxCyte Inc., USA). For each condition (WT and mutants), 25 µg of plasmid per transfection was used. Twenty-four hours after transfection, cells were dissociated with Accutase, diluted and transferred into the patch-clamp apparatus.

### Automated patch clamp (APC)

APC recordings were performed using the SyncroPatch 384PE from Nanion (Munich, Germany). Whole-cell T-type currents were recorded in transiently transfected HEK-293 cells. Single-hole, 384-well recording chips were used and seeded with 300,000 cells/mL. Pulse generation and data collection were performed with the PatchControl384 v1.5.2 software (Nanion) and the Biomek v1.0 interface (Beckman Coulter). After initiating the experiment, cell catching, sealing, whole-cell formation, buffer exchanges, recording, and data acquisition were all performed sequentially and automatically. The recording solutions were purchased from Nanion. The intracellular solution contained (in mM): 10 CsCl, 110 CsF, 10 NaCl, 10 EGTA, and 10 HEPES (pH 7.2). The extracellular solution contained (in mM): 140 NaCl, 4 KCl, 1 MgCl_2_, 5 Glucose, 10 HEPES (pH 7.4) with the final concentration of CaCl_2_ adjusted to 5.2 mM. This concentration was chosen to obtain greater calcium current density and higher percentage of gigaseal recordings. Whole-cell experiments were performed at a holding potential of -100 mV at room temperature (23°C). Currents were sampled at 20 kHz. Activation and inactivation curves were built using depolarization steps lasting 3000 ms from −120 mV to +50 mV, with 5 mV increments followed by a 200 ms depolarization step to −20 mV. Deactivation curves were built with a 20 ms pulse to -20 mV followed by 100 ms hyperpolarizing pulses from -120 mV to -60 mV. The recovery from inactivation was investigated using a double pulse protocol. Cells were first depolarized by a 1000 ms pre-pulse at −20 mV followed by a 20 to 7000 ms long interpulse interval at holding potential (-100 mV), and finally depolarized by a 100 ms test pulse at −20 mV.

### Manual clamp (MPC) and action potential (AP) clamp

Two days after transfection, cells were split at low density for whole-cell calcium current recordings using the patch-clamp technique with an Axopatch 200B amplifier (Molecular Devices). Borosilicate glass patch pipettes were used with a resistance of 1.5 ∼ 2.5 MOhm when filled with an internal solution containing (in mM): 140 CsCl, 10 EGTA, 10 HEPES, and 3 CaCl_2_ (pH adjusted to 7.25 with NaOH, ~315 mOsm). The extracellular solution contained (in mM): 135 NaCl, 20 TEACl, 2CaCl_2_, 1 MgCl_2_ and 10 HEPES (pH adjusted to 7.25 with NaOH 1M, ~330 mOsm). Recordings were filtered at 5 kHz. For the action-potential clamp studies performed in MPC, the stimulation commands were (1) a regular train of spikes recorded in Purkinje neurons of the cerebellum generously provided by Dr B. P. Bean (Harvard Medical School, Boston, MA, USA) (Raman and Bean, 1997), and (2) a reticular thalamic neuron (nRT) rebound burst (Chemin et al., 2002).

### Neuronal *in silico* modeling

Modeling was performed using the NEURON simulation environment (Hines and Carnevale, 1997). The model of cerebellar nuclear neuron is based on a previously published model (Sudhakar et al., 2015), downloaded from the NEURON database at Yale University (https://modeldb.science/185513)]. Neuronal activities were generated using the medium value of input gain, as described previously (Sudhakar et al., 2015). The electrophysiological properties of the Ca_v_3.1 channels were modeled using Hodgkin-Huxley equations as described previously (Huguenard and Prince, 1992;Destexhe et al., 1996). The values obtained for the Ca_v_3.1 WT and the variant channels were substituted for the corresponding values of native T-type channels in cerebellar nuclear neurons after fitting them with the initial model values in GraphPad Prism (see equations below). The membrane voltage values were corrected for liquid junction potential, which was 4.5 mV in the recording conditions.

**Activation steady state (minf)** = 1.0 / (1 + exp((v - v_1/2__minf)/k_minf))

**Inactivation steady state (hinf)** = 1.0 / (1 + exp((v - v_1/2__hinf)/k_hinf)

**Activation kinetics (taum)** = (C_taum + 0.333 / (exp((v - v_1/2__taum1)/ k_taum1) + exp((v - v_1/2__tau_m2)/k_taum2)))

**Inactivation kinetics (tauh)** = (C_tauh + 0.333 / exp((v - v_1/2__tauh1)/k_tauh1))

### Statistical analyses

Data were analyzed with GraphPad Prism and results are presented as means ± standard error of the mean (SEM). P-values for the statistical analyses were calculated using nonparametric (Kruskal-Wallis) one-way ANOVA followed by Dunnett’s post hoc multiple comparison test with the following significance criteria * *p*<0.05, ** *p*<0.01, and *** *p*<0.001.

## Results

### Current density measurements in APC

The 18 Ca_v_3.1 variants that we transiently transfected for electrophysiological characterization using APC are presented in **Table 1** and in **Figure 1**. Calcium currents were recorded using MPC for 10 of these variants, p.M197R, p.L208P, p.V392M, p.F956del, p.A961T, p.I962N, p.I1412T, p.M1531V, p.G1534D, and p.R1715H, (Coutelier et al., 2015;Chemin et al., 2018;Berecki et al., 2020;Qebibo et al., 2024). Eight variants have not yet been characterized: p.R102Q, p.V184G, p.N1200S, p.S1263A, p.R1718G, p.R1813W, p.V1835M, and p.D2242N. Among them, the p.S1263A and p.R1718G variants were recently reported (Riquet et al., 2023;Qebibo et al., 2024) but not yet electrophysiologically characterized. Variants located on the S4 segments are shown in green, those on the S5 segments in blue, the S6 segments in red, and those on the loops in grey (**Figure 1A**) with 15 of them being positioned in the cryo-EM structure of the Cav3.1 protein (**Figure 1B** ; (Zhao et al., 2019)). In APC experiments, the ratio of Ca_v_3.1 current-positive cells was in the range of 75% (comparing Ca_v_3.1 WT transfected cells with mock transfected cells). Only cells with a current density greater than 5pA/pF were considered for further detailed calcium current analyses. This led us identify 6 variants (p.M197R, p.V392M, p.F956del, p.I962N, p.I1412T, and p.G1534D) as showing current densities significantly lower than Ca_v_3.1 WT channels (**Figure 2A**). Four of these variants, p.M197R, p.F956del, p.I1412T, and p.G1534D, previously characterized using MPC (Qebibo et al., 2024), resulted in too few cells with large enough calcium current density for accurate biophysical characterization in APC experiments. These 4 variants, as well as p.V392M variant for which only half-activation potentials could be determined, were not considered for further analysis of their APC recordings. Only the p.I962N variant exhibited a sufficient average current density and quality traces for complete characterization via APC. Contrasting with low expressing variants, p.V184G, p.N1200S, p.S1263A and p.D2242N variants displayed significantly higher current densities (**Figure 2A**). The superimposed representative current traces at -20mV (**Figure 2B-E**) illustrate the variations in current density for all these variants, especially the ones with nearly null current density (p.M197R and p.I1412T). In addition, the current traces show the marked differences in inactivation kinetics especially for most of the variants located on the S5 and S6 segments defining the IG (**Figure 2C-D**).

**Table 1:** Presentation of the 18 *CACNA1G* variants investigated in the study in automated patch-clamp (APC), manual patch-clamp (MPC) or both (APC / MPC). The clinical description of the variants in black can be found in previous studies. The variants in bold-red are reported for the first time, with a brief clinical description of the related patients. Bold-underlined MPC indicates that the properties of these variants were preferentially obtained in MPC.

| Variants | MPC / APC tested | Clinical information |
| --- | --- | --- |
| **p.R102Q** | APC | Ataxia, progressive cerebellar atrophy, global developmental delay. Medical history complicated by prenatal exposures to drugs/alcohol. Possible encephalitis in infancy. Variant also present in EXAC. |
| **p.V184G** | APC | Adult-onset neuromuscular disease, including ptosis, muscle weakness, peripheral neuropathy, and ataxia. |
| p.M197R | APC / **MPC** | (Qebibo et al., 2024) |
| p.L208P | APC | (Berecki et al., 2020) |
| p.V392M | APC / MPC | (Qebibo et al., 2024) |
| p.F956del | APC / **MPC** | (Qebibo et al., 2024) |
| p.A961T | APC / MPC | (Chemin et al., 2018;Qebibo et al., 2024) |
| p.I962N | APC / MPC | (Qebibo et al., 2024) |
| **p.N1200S** | APC / MPC | (Kosmicki et al., 2017) |
| p.S1263A | APC | (Qebibo et al., 2024) |
| p.I1412T | APC / **MPC** | (Qebibo et al., 2024) |
| p.M1531V | APC / MPC | (Chemin et al., 2018;Qebibo et al., 2024) |
| p.G1534D | APC / **MPC** | (Qebibo et al., 2024) |
| p.R1715H | APC / MPC | (Coutelier et al., 2015) |
| p.R1718G | APC | (Riquet et al., 2023;Qebibo et al., 2024) |
| **p.R1813W** | APC | Congenital ataxia. Found in one patient, inherited from his mother, both also having a known pathogenic *CACNA1A* variant but with incomplete penetrance/expressivity. |
| **p.V1835M** | APC / MPC | A 2.5-year-old girl (in 2019) with developmental delay, microcephaly, and tremor (when scared/anxious). Exome sequencing revealed a pathogenic variant in *MECP2*, so she has Rett syndrome. She also has tremor. Her brain MRI at 1.5 years was normal; it did not show cerebellar atrophy. |
| **p.D2242N** | APC | Mild intellectual disability, ophthalmoplegia, and progressive ataxia, dysarthria, and dysphagia. He has developed prognathism. Brain MRI shows olivopontocerebellar atrophy |

**Figure 1:**
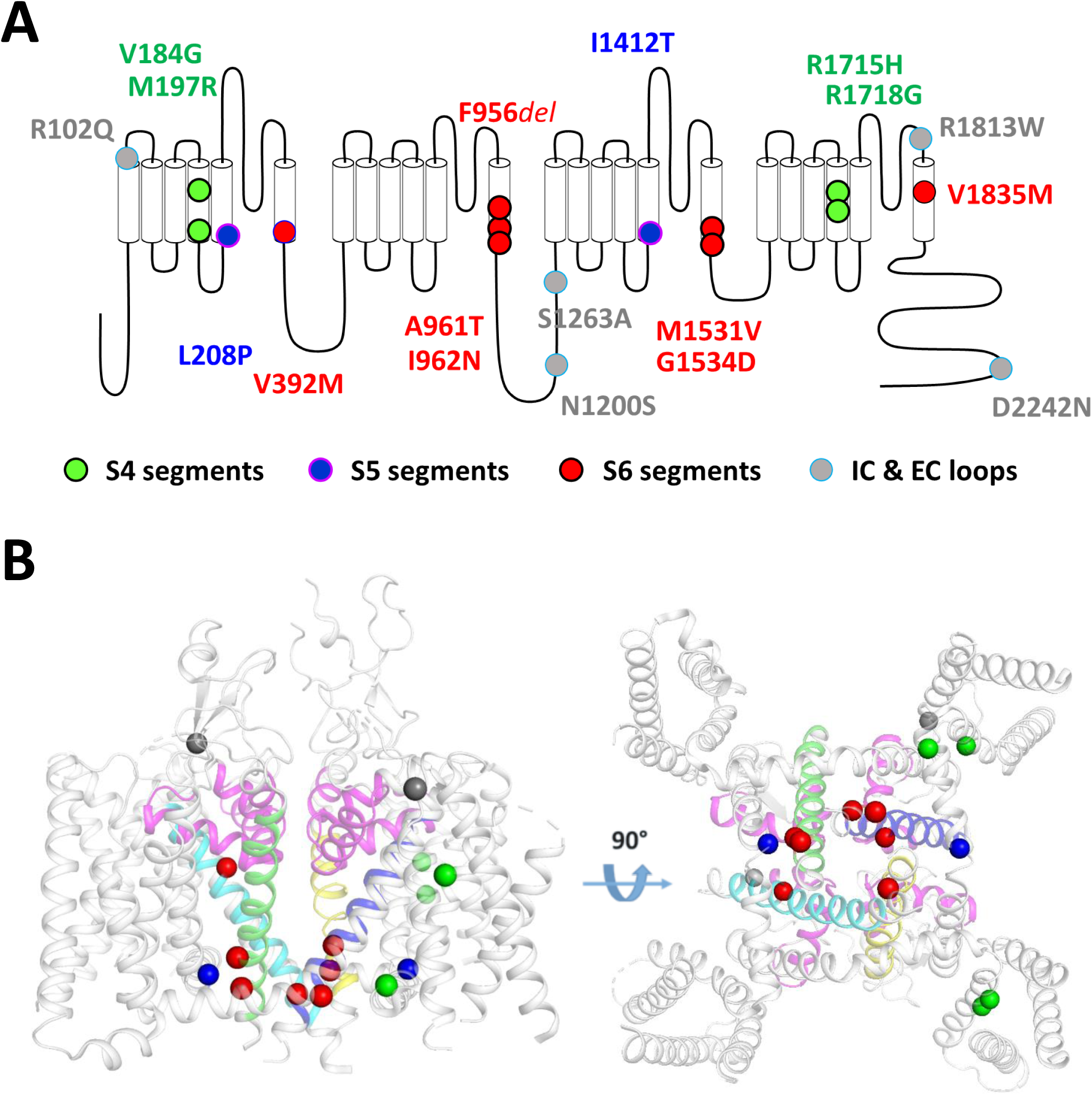
Schematic localization on the Cav3.1 channel of the 18 variants investigated in this study. **(A)** The transmembrane topology of the Cav3.1 calcium channel shows the four domains repeat (DI to DIV), each composed of six transmembrane segments (S1 to S6). Mutations on segments S4, S5, S6, and the intracellular and extracellular loops (IC, EC) are depicted in green, blue, red, and gray, respectively. **(B)** 3D representation of the position of 15 variants on the Cryo-EM resolved structure of Cav3.1 (PDB: 6KZO ; (Zhao et al., 2019)) with a side view (left panel) and a bottom view (right panel). The 3 additional variants (p.N1200S, p.S1263A and p.D2242N) are found in area of the Cav3.1 protein that were not resolved in the 6KZO PDB structure.

**Figure 2:**
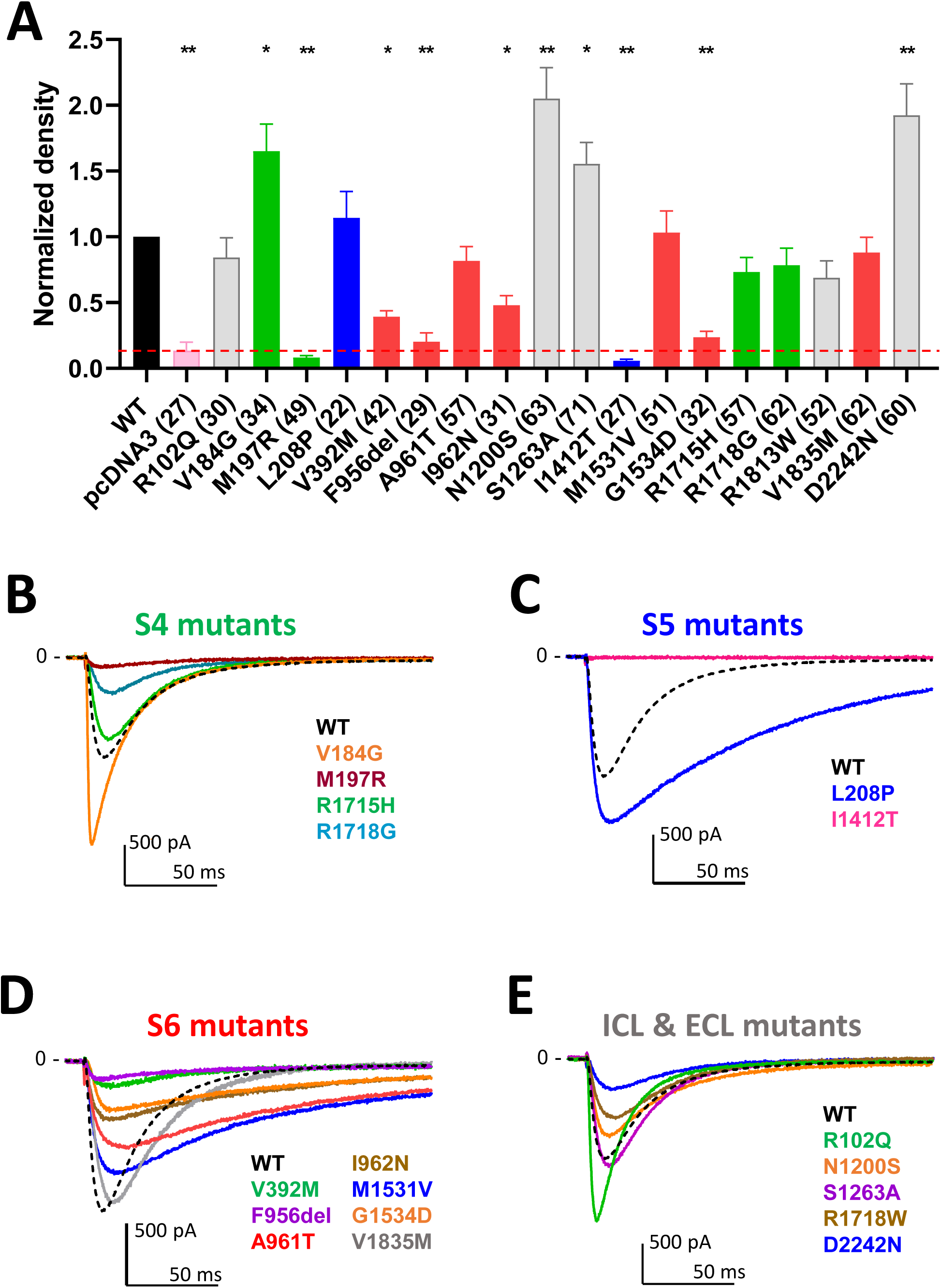
Density and calcium current traces for the 18 Cav3.1 variants studies in APC. **(A)** Graph of the calcium current density at -20 mV for all the variants, normalized to the WT current density. The red dotted line indicates the maximum density obtained on empty pcDNA3-transfected cells (∼ 5pA/pF) that was used to identify low-expressing cells excluded for further biophysical analysis. **B - E.** Current trace examples at -20 mV for the all the variants grouped according to their localization in the Cav3.1 protein. Distinct colors were used to better visualize the calcium current traces illustrating each variant of the S4 group (**B**), the S5 group (**C**), the S6 group (**D**) and the group comprising variants in loops, N-ter and C-ter (**E**), compared to WT (black dotted line).

### Biophysical properties of variants characterized by APC

The activation kinetics is rather fast for the WT Ca_v_3.1 current (∼ 4ms at -20mV) in APC recordings. The S4 variants p.R1715H and p.R1718G, as well as the S6 variant p.V1835M, exhibit slower activation, while most of the variants in S5 and S6 segments display faster activation (**Figure 3A** and **Supplementary Figure 1**), compared to WT. Similar to that reported in MPC, the inactivation and deactivation kinetics are significantly slower for the S5 and S6 IG variants (p.L208P, p.A961T, p.I962N, p.M1531V) (**Figure 3B-C**, and **Supplementary Figures 2 and 3**). Regarding recovery from inactivation, the p.L208P, p.A961T, and p.I962N variants are the slowest and do not fully recover from inactivation (**Figure 3D** and **Supplementary Figure 4**), again in good agreement with MPC experiments. Notably, all the variants evaluated in APC experiments display no shift or a negative shift in their steady-state activation and inactivation properties (**Figure 4A-B**). Similar to that reported in MPC experiments, these hyperpolarizing shifts were highly significant for the IG variants p.L208P, p.V392M, p.A961T, p.I962N and p.M1531V.

**Figure 3:**
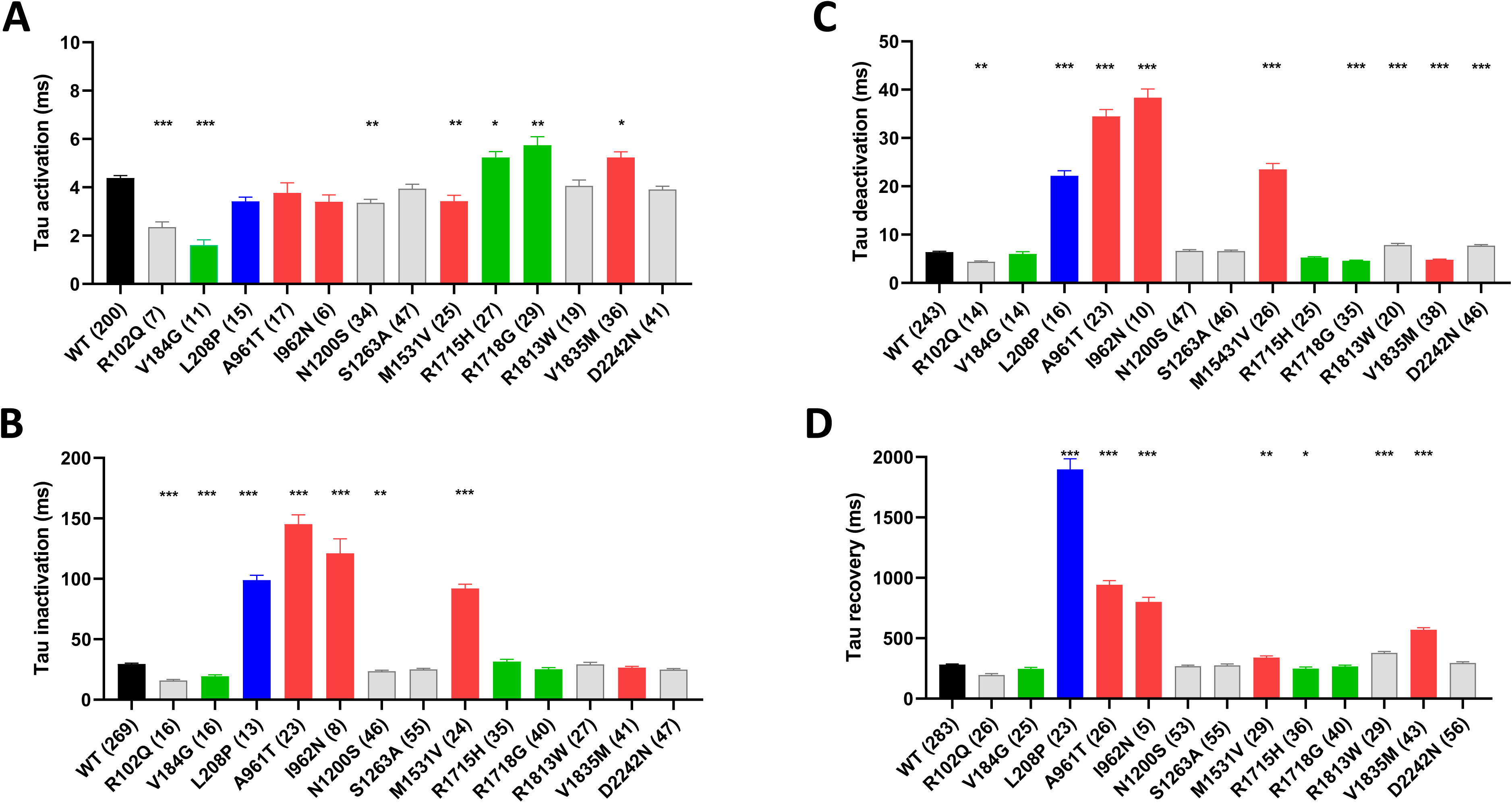
Kinetic properties of Cav3.1 variants using APC. **(A)** Activation kinetics at -20 mV. (**B)** Inactivation kinetics at -20 mV. **(C)** Deactivation kinetics at -60 mV. **(D)** Recovery from inactivation. Note that the recordings for the variants p.M197R, p.V392M, p.F956del, p.I1412T and p.G1534D did not meet quality controls for measuring all the electrophysiological parameters.

**Figure 4:**
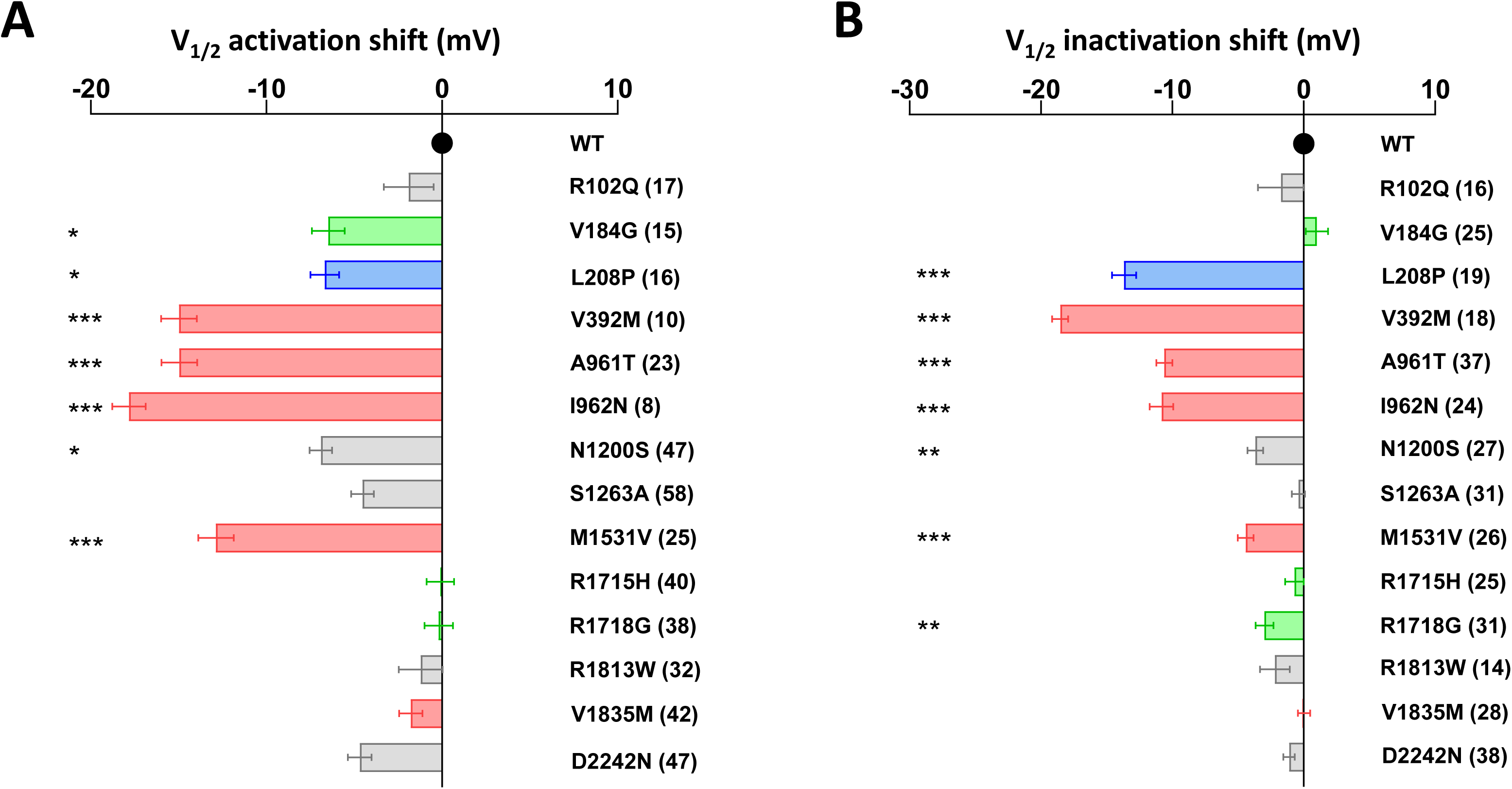
Half-activation and inactivation properties of Cav3.1 variants using APC. **(A)** The shift in steady-state half-activation potential (V_1/2_ activation) for each variant is compared to the WT value. **(B)** The shift in steady-state half-inactivation potential (V_1/2_ inactivation) for each variant is compared to the WT value.

### Correlation between APC and MPC data

Superimposed current-voltage relationships (**Figure 5A**), steady-state activation (**Figure 5B**) and steady-state inactivation (**Figure 5C**) recorded in MPC and APC for the WT and the most recurrent SCA42ND variant (p.A961T) illustrate well that the hyperpolarizing shift in steady-state properties for the p.A961T variant was comparable in MPC and APC experiments. The use of a higher extracellular calcium concentration in APC (5 mM), compared to MPC (2 mM), was responsible for a depolarizing shift of similar amplitude for both WT and p.A961T variant in the steady-state activation (15 mV for WT and 16 mV for p.A961T) and in the steady-state inactivation (4 mV for WT and 3 mV for p.A961T). We then assessed the correlation between the electrophysiological properties in APC and MPC for the set of variants studied in both experiments (**Figure 5D-G**). Indeed, both the steady-state activation (V_1/2act_) and inactivation (V_1/2inact_) showed a robust correlation (r close to 1 and p<0.05) between APC and MPC (**Figure 5D-E**), as well as for deactivation kinetics and recovery from inactivation (**Figure 5G** and **Supplementary Figure 5**). Correlation was however not observed for the inactivation kinetics data (r = 0.66), likely due to the dispersion of values for slow inactivating mutants, compared to the highly clustered WT-like mutants (**Figure 5F**). This was similarly true for the activation kinetics (**Supplementary Figure 5**).

**Figure 5:**
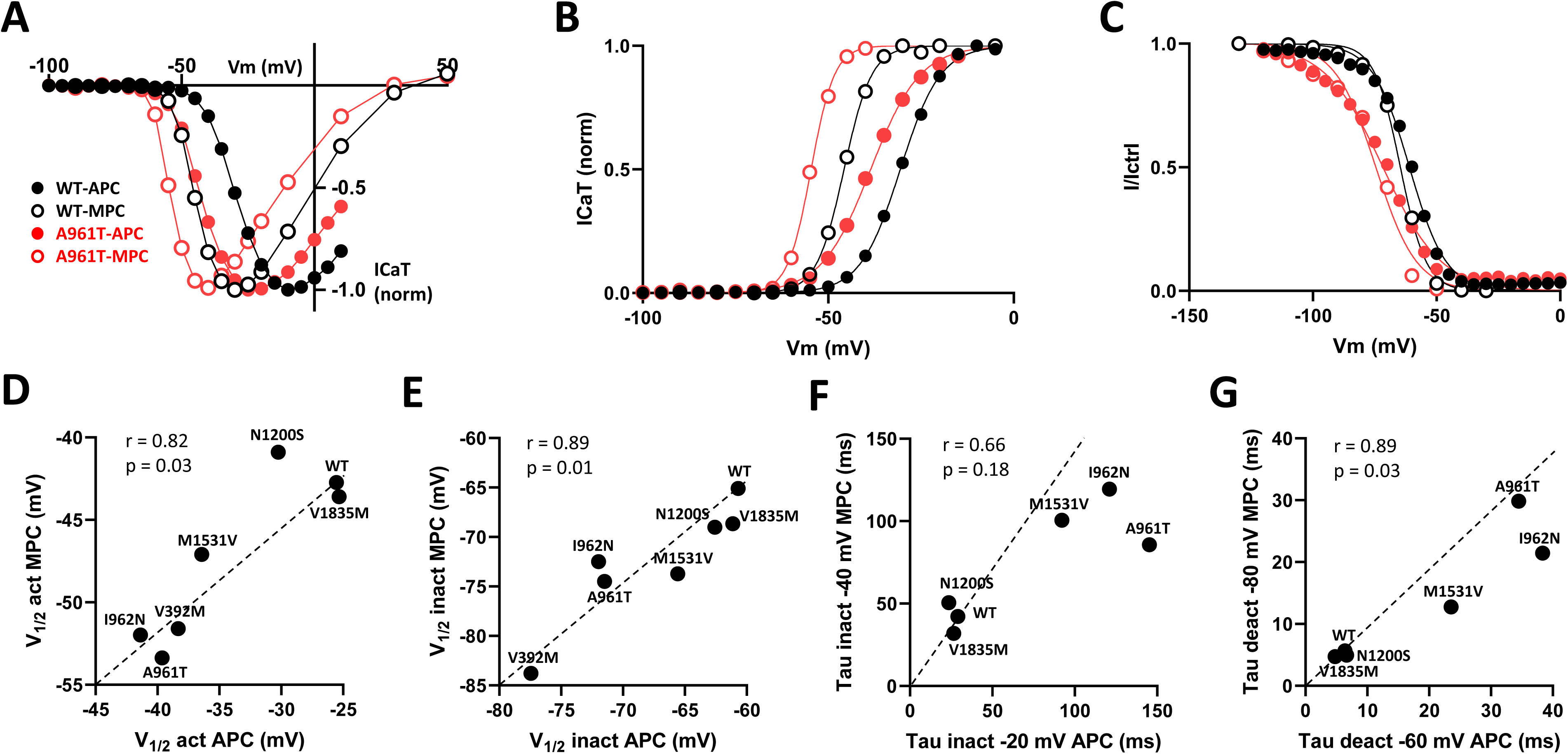
Comparison of MPC and APC electrophysiological parameters. **(A)** I-V curves obtained using MPC (open symbols) and APC (filled symbols) for WT (black) and the recurrent SCA42ND variant p.A961T (red). **(B)** Steady-state activation curves. **(C)** Steady-state inactivation curves. **(D** to **G).** The correlation graphs for half-activation potential **(D)**, half-inactivation potential **(E)**, inactivation kinetics **(F)** and deactivation kinetics **(G)**, respectively, measured in APC (X axis) and MPC (Y axis).

### Predicted consequence of Ca_v_3.1 variants properties on neuronal excitability

The electrophysiological parameters collected in MPC and APC experiments (V_1/2_ act, V_1/2_ inact, Kact, Kinact, Tau act, Tau inact, Tau deact) can be used in a virtual model of DCN (Sudhakar et al., 2015) neurons to estimate the variants’ effect on DCN firing activity by measuring the action potential (AP) frequency (**Figure 6, Supplementary Figures 6-8**). We recently reported that the IG variants investigated in MPC (p.V392M, p.F956del, p.A961T, p.I962N, p.I1412T, p.M1531V, p.G1534D) produced a higher AP frequency compared to WT (Qebibo et al., 2024). It is notable that with the parameters of the variants fully explored with the APC approach, the p.A961T, p.I962N, p.M1531V variants, and to a lesser extend the p.L208P variant, also showed a higher AP frequency, pointing further to the GOF properties of these variants (**Figure 6A**) as exemplified for p.A961T compared to WT (**Figure 6B-C**). Because APC experiments have revealed that Ca_v_3.1 variants exhibit either higher or lower current density compared to WT channel (**Figure 2A**), we next studied the consequence of varying the current density on AP-frequency for all the variants studied in APC (**Figure 6D**) and all the variants studied in MPC (**Figure 6E**). A 2-fold increase in current density led to a marked increase in AP-frequency for the GOF variants, especially p.L208P (**Figure 6D**). WT-like variants also displayed increased AP-frequency but was unchanged for the loss-of-channel-activity variant p.M197R (**Figure 6E**). Finally, when the DCN modeling was performed using the Ca_v_3.1 current density corresponding to that measured in APC (see **Figure 2A**), only the IG variants p.M1531V > p.A961T > p.L208P > p.I962N, as well as the variants p.V184G and p.N1200S showed increased AP-frequency (**Figure 6F-G and Supplementary Figures 7-8**).

**Figure 6:**
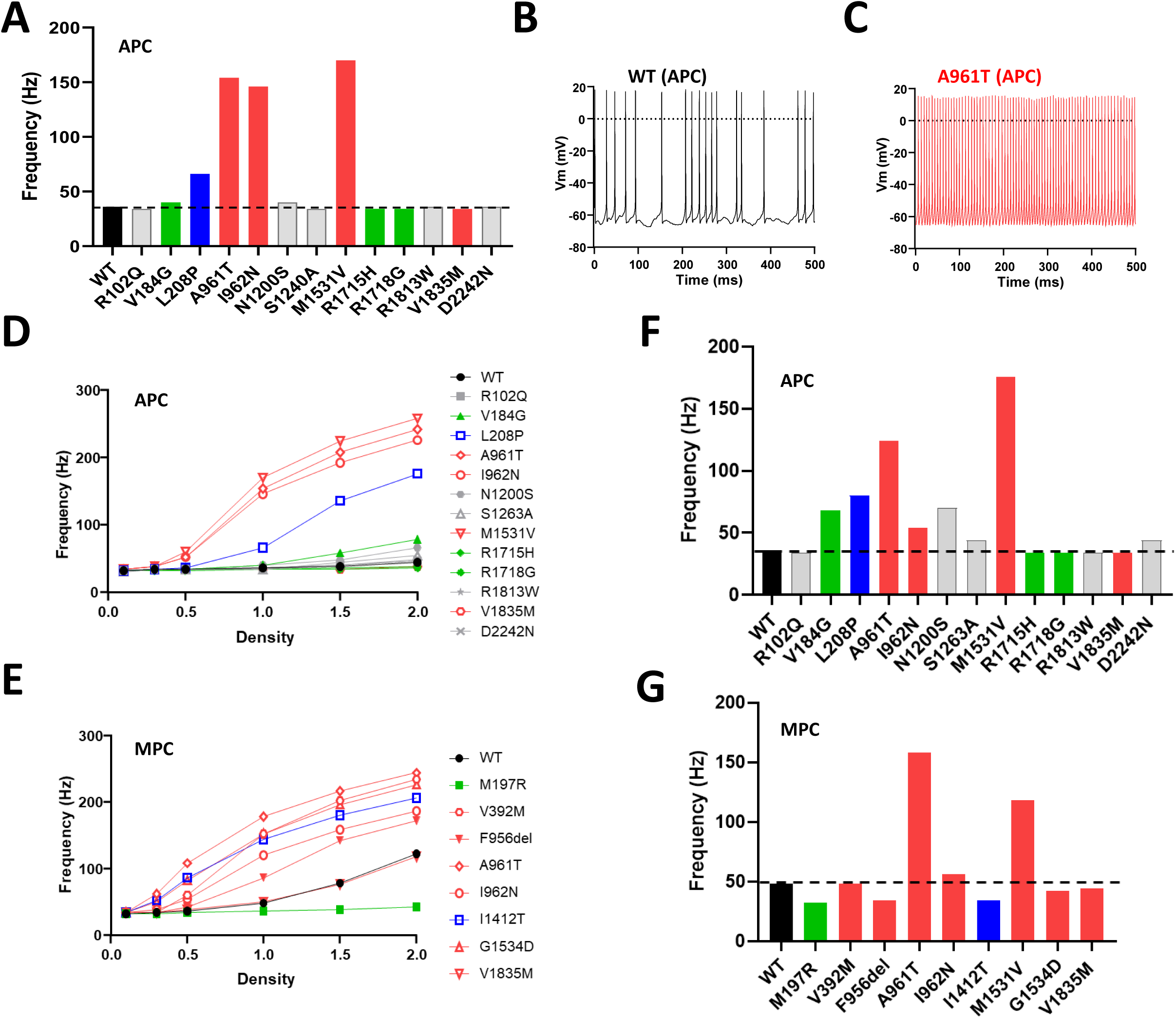
Deep Cerebellar Nucleus (DCN) modeling using Cav3.1 variant parameters obtained in APC and in MPC experiments. **(A)** Spike frequency graph for Cav3.1 variants characterized using APC. **(B** and **C)** Examples of DCN spiking activity for using APC parameters obtained for WT **(B)** and p.A961T channels **(C)**. **(D** and **E)** Change in DCN spike frequency obtained with increasing current densities for all the variants characterized in APC **(D)**, and in MPC **(E)**, respectively. **(F** and **G)** The DCN spike frequency obtained for the current density measured in Figure 2B (normalized to the WT current density) for the variants characterized in APC **(F)** and MPC **(G)**, respectively.

### GOF effect validation in action-potential clamp experiments for p.M1531V and p.A961T

A further functional evaluation of the GOF effect of four representative IG variants (p.V392M, p.A961T, p.M1531V, p.G1534D) was performed using action potential (AP) clamp experiments in MPC (**Figure 7**). The calcium current in HEK-293 cells expressing p.V392M, p.A961T, p.M1531V or p.G1534D variants was recorded during an AP-voltage command, mimicking (i) a tonic firing activity of Purkinje cells and (ii) a rebound burst firing from thalamic nRT neurons. In these recordings, the resulting calcium current accounts for the specific electrophysiological behavior of each variant (**Figure 7A-B**). For the tonic firing activity, it is worth noting that most of the inward current occurred during the interspike, in link with the slow deactivation of Ca_v_3.1, even slower for IG variants (**Figure 7A**). For the rebound burst firing activity, the calcium current relies both on Ca_v_3.1 de-inactivation and slow deactivation (**Figure 7B**). The area under the curve was measured and normalized to the maximum amplitude of the current density measured in each cell using a standard test-pulse protocol. The p.A961T and p.M1531V variants showed a significant increase in calcium current entry in both tonic and rebound burst activities (**Figure 7C-D**). On the contrary, the p.V392M and p.G1534D variants showed a reduced calcium current during the tonic firing activity (**Figure 7C**). Using rebound burst activities, the p.G1534D variant exhibited a moderate increase in calcium current, while the p.V392M variant showed a lower calcium entry, compared to WT (**Figure 7D**). These AP clamp experiments highlight that the Ca_v_3.1 variant-dependent calcium entry relies on the electrophysiological behavior of neurons and further confirm the GOF properties of the p.A961T and p.M1531V variants.

**Figure 7:**
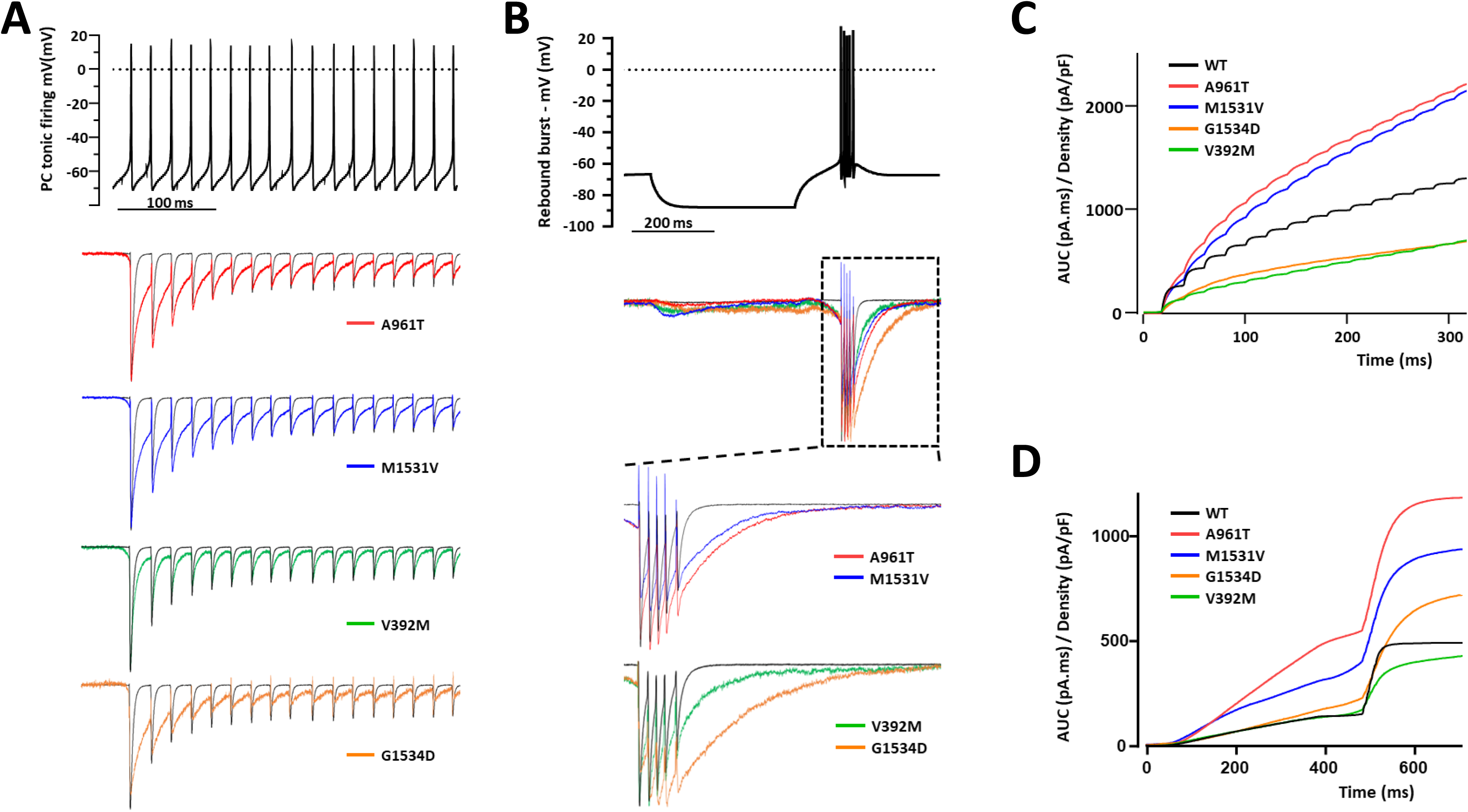
Action-potential clamp experiments with IG variants. **(A)** Representative calcium current traces for p.A961T, p.M1531V, p.V392M, and p.G1534D variants (lower panel) in response to a Purkinje cell tonic firing activity used as voltage command (upper panel). (**B)** Representative calcium current traces for p.A961T, p.M1531V, p.V392M, and p.G1534D variants (lower panel) in response to a thalamic rebound burst firing activity used as voltage command (upper panel). (**C** and **D)** The integral (area under the curve) quantification of the calcium current for each variant obtained for the tonic firing activity (**C**) and the rebound burst activity (**D**), respectively.

## Discussion

### Automated patch-clamp techniques for classification of the Cav3 channelopathies

The APC technology is revolutionizing the field of electrophysiology, particularly for the study of channelopathies, by allowing investigation of a large number of disease-related variants associated with various neurological and cardiac disorders (Vanoye et al., 2021). APC allows to record and analyze hundreds of cells in parallel, for a more comprehensive characterization of ion channel variants (Ng et al., 2021;Ma et al., 2023). Traditional MPC techniques are highly accurate in describing parameters, but time-consuming and labor-intensive, while APC offers high-throughput capability and may be less operator-dependent (Yajuan et al., 2012). This is particularly crucial given the growing number of identified ion channel variants (Lukacs et al., 2021;Obergrussberger et al., 2022) as here with the discovery of many novel *CACNA1G* variants associated to neurological conditions. We and others have shown that APC experiments using the SyncroPatch PE384 are reliable in transposing electrophysiological parameters to study voltage-gated sodium and potassium channels, provided that appropriate guidelines are respected (Glazer et al., 2020;Montnach et al., 2021;Oliveira-Mendes et al., 2023). T-type Ca_v_3 calcium channels are particularly well suited for APC investigations as only the pore channel protein (Ca_v_α_1_) is required to produce a native-like T-type calcium current (Chemin et al., 2002;Perez-Reyes, 2003). Indeed, APC was recently used to study 57 variants of the Ca_v_3.3 channel identified in a large schizophrenia cohort (Baez-Nieto et al., 2022). Here we have also successfully used APC to characterize 18 Ca_v_3.1 disease variants, especially 8 variants of uncertain significance (VUS) that are studied at the functional level for the first time. Our study validates that MPC experimental conditions could be adequately transposed to APC for Ca_v_3.1 study. Importantly, the gating defects observed for the *de novo* IG variants (S5-S6 segments) using MPC i.e., slow inactivation and deactivation kinetics, hyperpolarizing shift of steady-state activation and inactivation properties (Chemin et al., 2018;Berecki et al., 2020;Qebibo et al., 2024) were accurately replicated in APC experiments. This was also validated for the other variants studied in both MPC and APC experiments. APC offers to record a large number of cells, and blindly, for each condition (variant), which is well suited for measuring the variant current density. While some variants displayed increased current density, our study also points to a few variants showing very low current density. According to our quality control criteria, low expressing (LOF) variants resulted in a small number of cells that could be accurately studied in APC. The difficulty to study LOF variants may be a caveat when examining large series disease variants using APC only. Overall, the APC and MPC approaches were highly complementary in providing a comprehensive electrophysiological analysis of our large series of Ca_v_3.1 variants.

### Deciphering the gain/loss of channel activity in support of GOF or LOF variants

Several GOF variants in *CACNA1G* (*de novo*, missense mutations) are now linked to a variety of neurological and neurodevelopmental diseases with some severe conditions as SCA42ND (Chemin et al., 2018;Qebibo et al., 2024). Deciphering the electrophysiological alterations caused by these mutations to the Ca_v_3.1 channels is necessary to better document the disease mechanism(s), and to identify potential therapeutic opportunities. The electrophysiological criteria supporting a gain of channel activity are the increase in current density, the hyperpolarizing shift of the steady-state activation curve, the slower inactivation and deactivation kinetics and the increased window current (Chemin et al., 2018;Berecki et al., 2020;Qebibo et al., 2024). In turn, lower current density, slower recovery from inactivation and hyperpolarized steady state inactivation curve are in favor of loss of channel activity (Chemin et al., 2018;Qebibo et al., 2024). Most analyses of the channel variants’ gain/loss of channel activity are to date performed in heterologous expression systems, e.g. in transfected HEK-293 cells (as here), without considering some specificities of the native distribution of the studied channel. Recently, neuronal modeling was used to support MPC findings for several Cav3 variants and mutants (Blesneac et al., 2015;Coutelier et al., 2015;Chemin et al., 2018;El Ghaleb et al., 2021;Baez-Nieto et al., 2022;Qebibo et al., 2024). Ca_v_3.1 is highly expressed in several cerebellar neurons, especially in the deep cerebellar nucleus (DCN), for which virtual neuron models have been developed (Destexhe et al., 1996;Anwar et al., 2012;Sudhakar et al., 2015). We show here that the use of computed neuronal excitability (Sudhakar et al., 2015) allowed us to pinpoint the gain / loss of channel property for the eighteen variants explored in this study either in MPC, in APC or both. Coupling APC and MPC data with *in silico* neuronal modeling appear as a robust and complementary method for classifying Ca_v_3.1 variants. This approach further described the two recurrent SCA42ND variants p.A961T and p.M1531V as GOF variants.

In this study, we also report that the ‘gain/loss of channel activity’ toolbox could be completed with action potential (AP) clamp experiments. AP clamp experiments were performed in HEK-293 cells with neuronal activities originating from cells known to express Ca_v_3.1 (cerebellar and thalamic neurons) as voltage command (Chemin et al., 2002). The GOF ability of the two SCA42ND variants p.A961T and p.M1531V was retrieved both using tonic firing (Purkinje neuron) and rebound burst firing (thalamic neuron). The two other IG variants examined here, p.V392M and p.G1534D, both displayed reduced activity with tonic firing voltage command, while p.G1534D, but not p.V392M, show increased calcium entry in rebound burst firing. Overall, *in silico* modeling and AP clamp experiments revealed that at the functional level, the GOF properties of the variants are intimately associated with the specific electrophysiological signature of cells expressing the Ca_v_3.1 channel. Our study demonstrates that these experiments add to the variant characterization pipeline by contributing to a better classification of channel variants. To characterize further the GOF / LOF properties of *CACNA1G* variants, *in vivo* studies will also be instrumental. However, it will require to develop the appropriate models (human iPSC-derived cellular models or animal models) to take into account the diversity of channel variants, while complying with the 3R rule (Diaz et al., 2020).

## Data availability statement

The raw data supporting the conclusions of this article will be made available by the authors.

## Author contributions

**AD:** Investigation, Methodology, Data curation, Formal Analysis, Writing–original draft, Writing–review and editing. **LC:** Investigation, Methodology, Data curation, Formal Analysis, Writing–review and editing. **BR:** Methodology, Data curation, Formal Analysis, Writing–review and editing. **NB:** Investigation, Methodology, Data curation, Formal Analysis, Writing–original draft, Writing–review. **JM:** Methodology, Data curation, Formal Analysis, Writing–review and editing. **LB:** Investigation. Validation, Writing–review and editing. **VC:** Investigation. Validation, Writing–review and editing. **NW:** Validation, Writing–review and editing. **SN:** Validation, Writing–review and editing. **AM:** Validation, Writing–review and editing. **MDW:** Conceptualization. Formal analysis. Methodology. Funding acquisition. Supervision. Validation, Writing–review and editing. **PL:** Conceptualization. Formal analysis. Methodology. Funding acquisition. Supervision. Validation, Writing–original draft, Writing–review and editing.

## Funding

This research was funded by Labex “Ion Channel Science and Therapeutics” (ICST) and Agence Nationale de la Recherche (ANR-11-LABX-0015) and ‘Connaître les Syndromes Cérébelleux’ (CSC) Association (to P.L and L.B).

## Conflict of interest

The authors declare that the research was conducted in the absence of any commercial or financial relationships that could be construed as a potential conflict of interest.

## Supplementary material

The Supplementary figures for this article can be found in the attached file.

## Acknowledgments

The authors thank Dr Angela Sun (University of Washington, Seattle, USA) and Dr Julia Rankin (Royal Devon and Exeter Hospital, Wonford, UK) and all the clinicians involved in recruiting the cases presented in this study.

---

**Supplementary Figure 1 :**
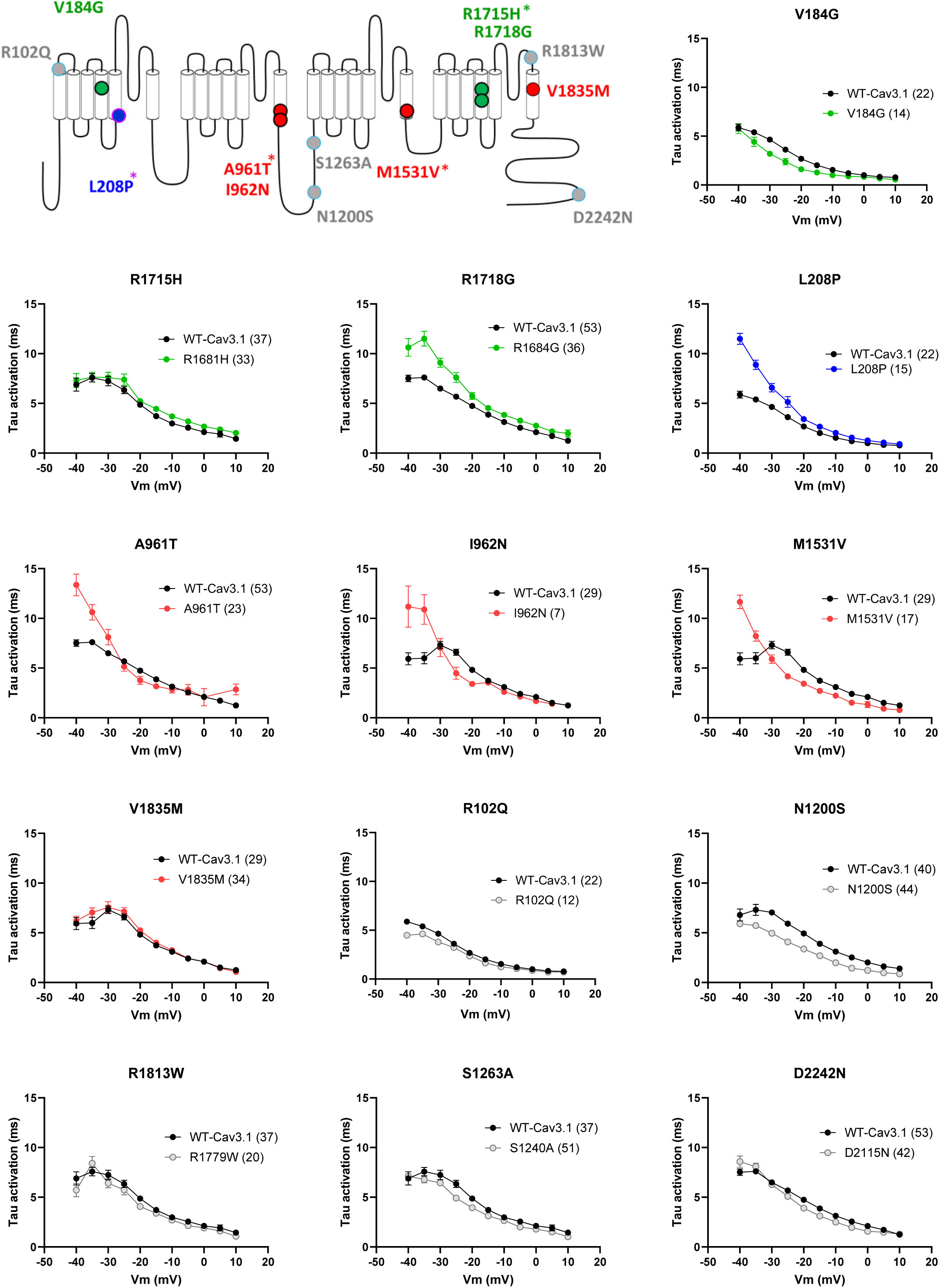
activation kinetics in APC.

**Supplementary Figure 2 :**
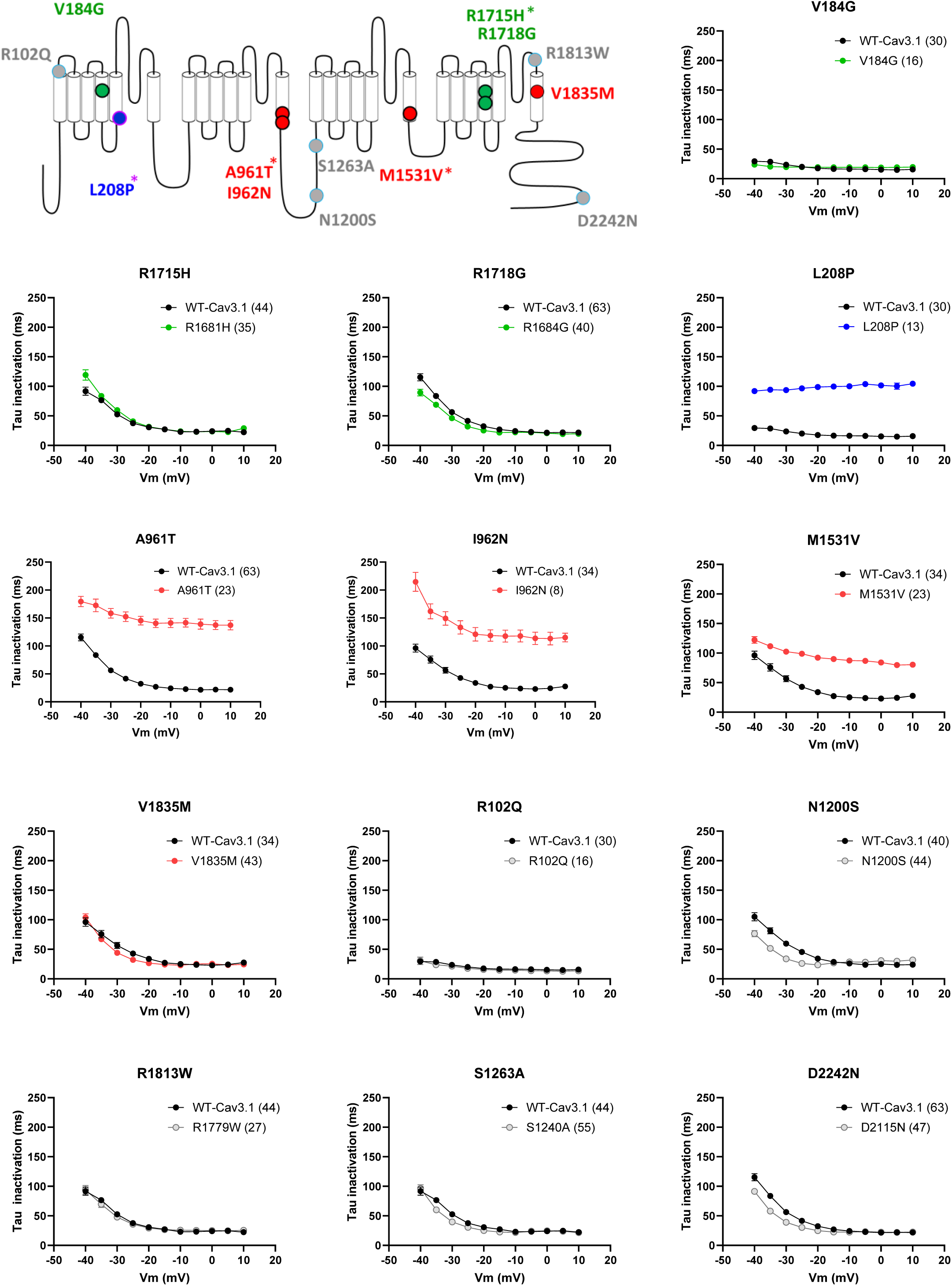
inactivation kinetics in APC.

**Supplementary Figure 3 :**
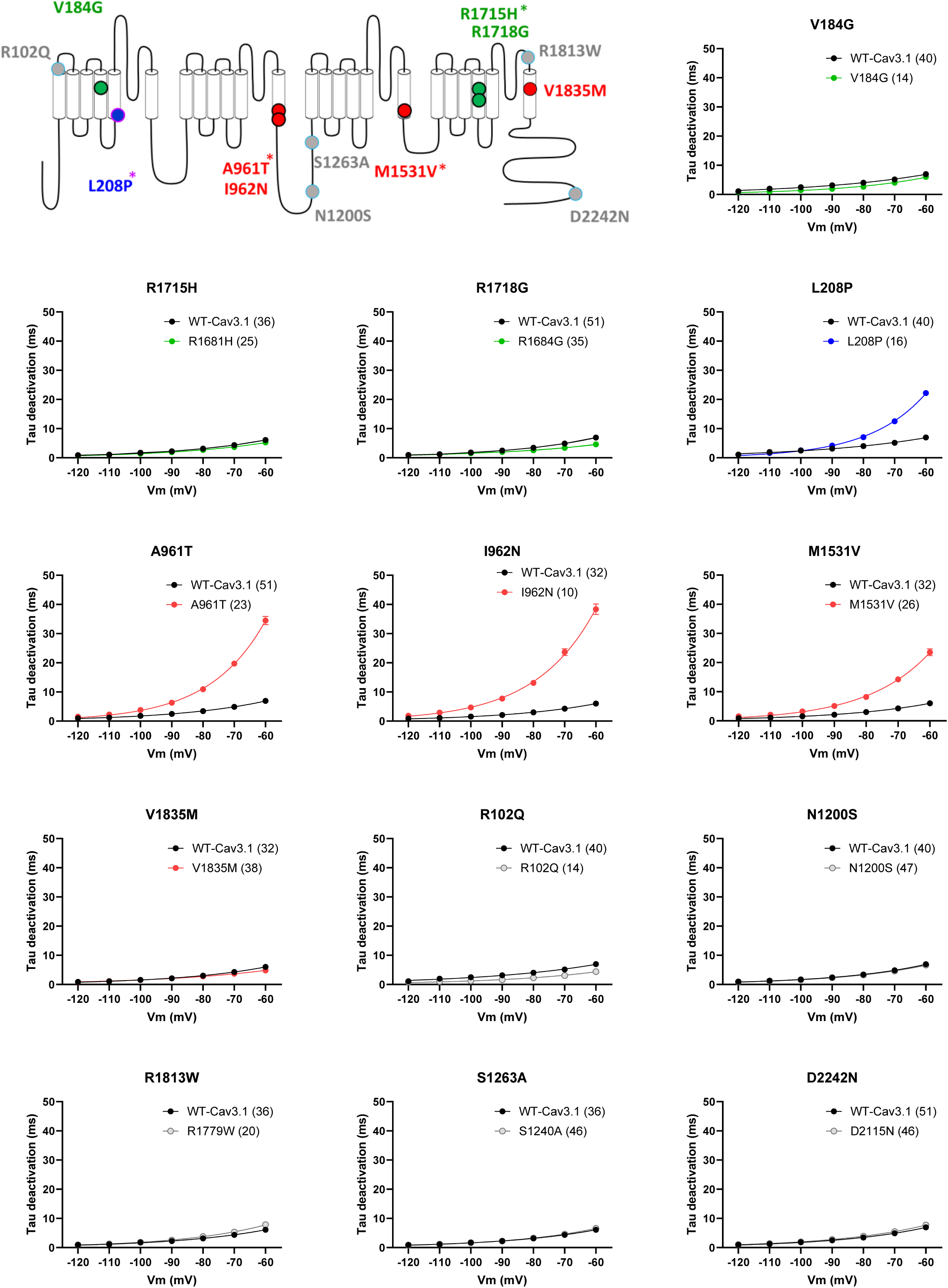
deactivation kinetics in APC.

**Supplementary Figure 4 :**
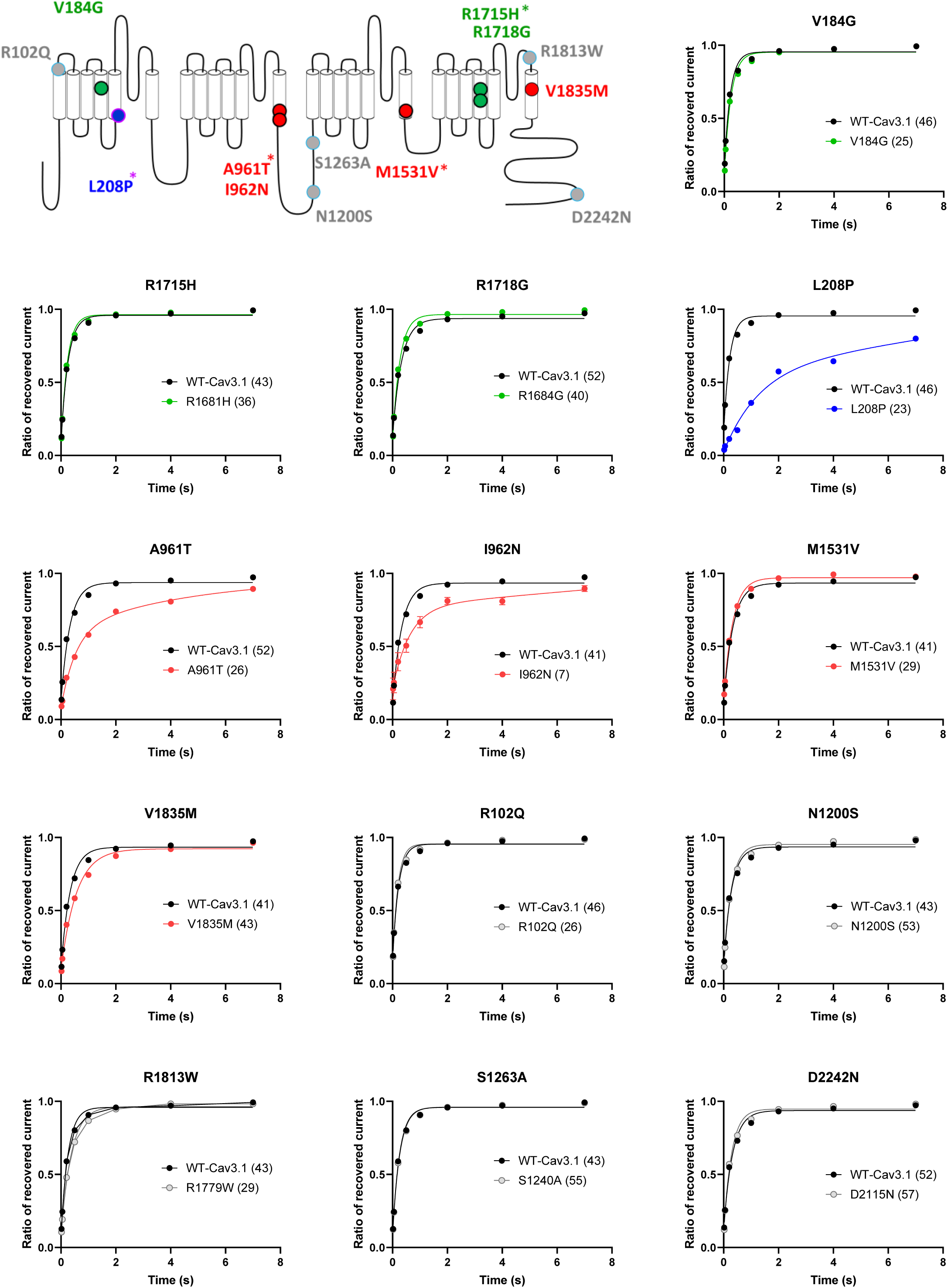
recovery from inactivation in APC.

**Supplementary Figure 5 :**
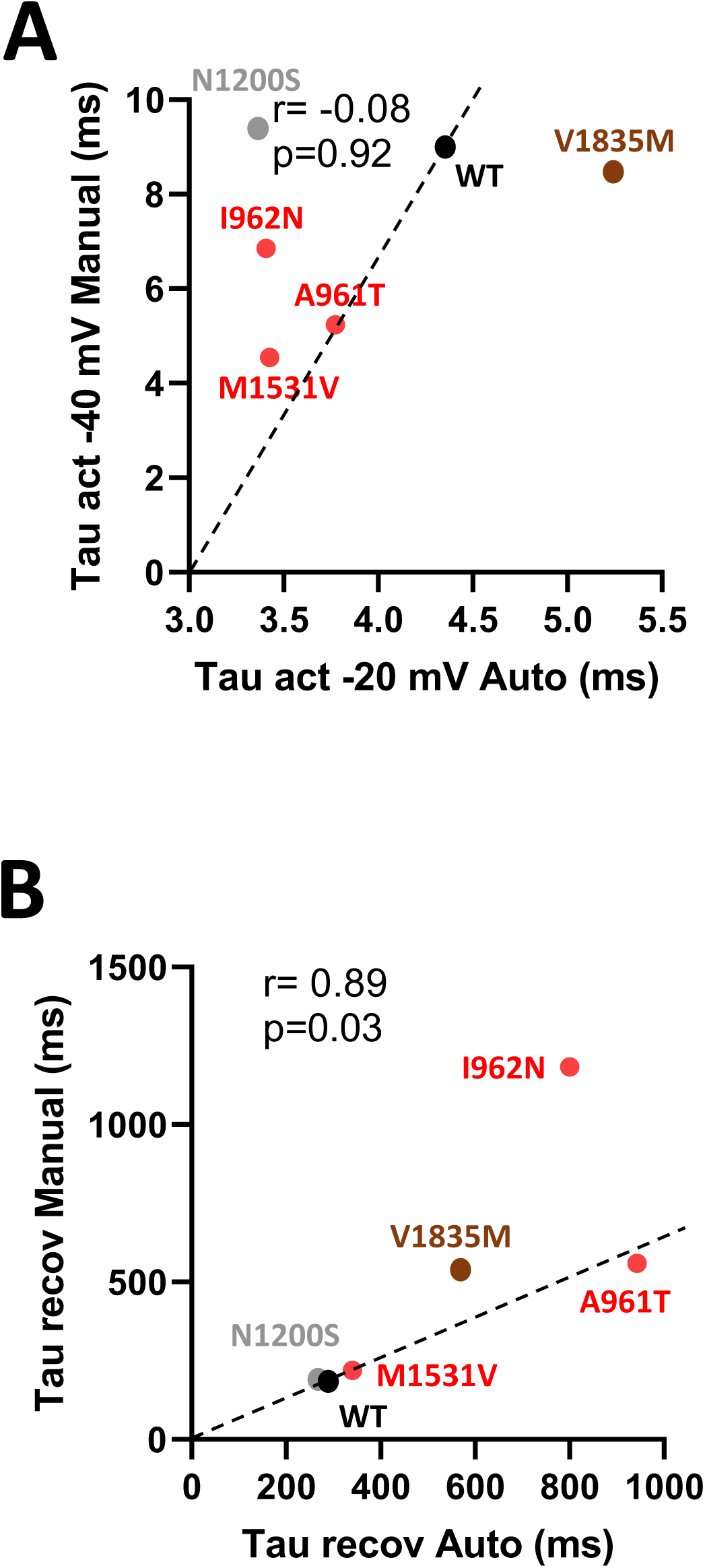
APC vs MPC correlation for activation kinetics (A) and recovery from inactivation (B)

**Supplementary Figure 6:**
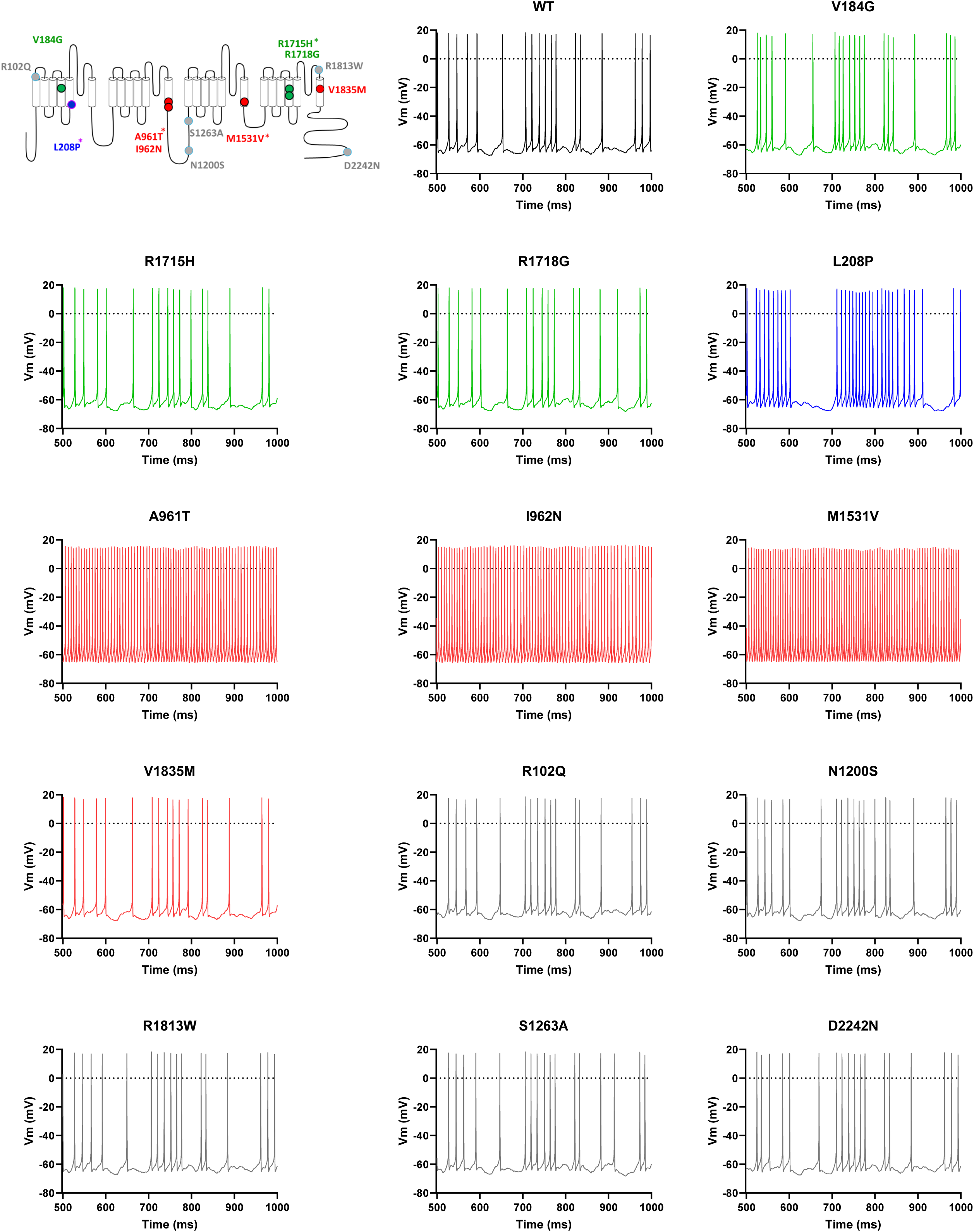
DCN modeling with APC electrophysiological parameters.

**Supplementary Figure 7:**
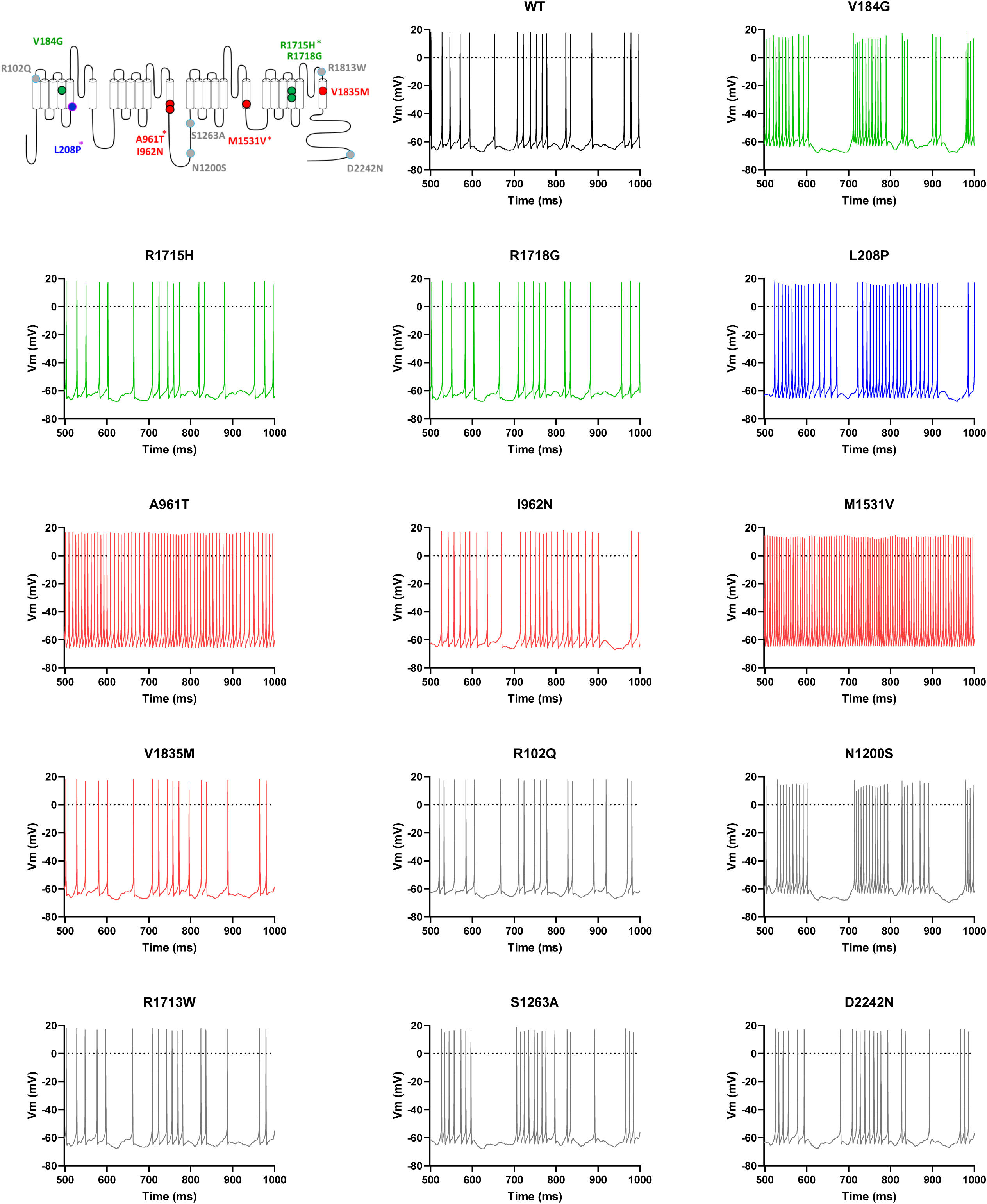
DCN modeling normalized for current density (APC param.)

**Supplementary Figure 8:**
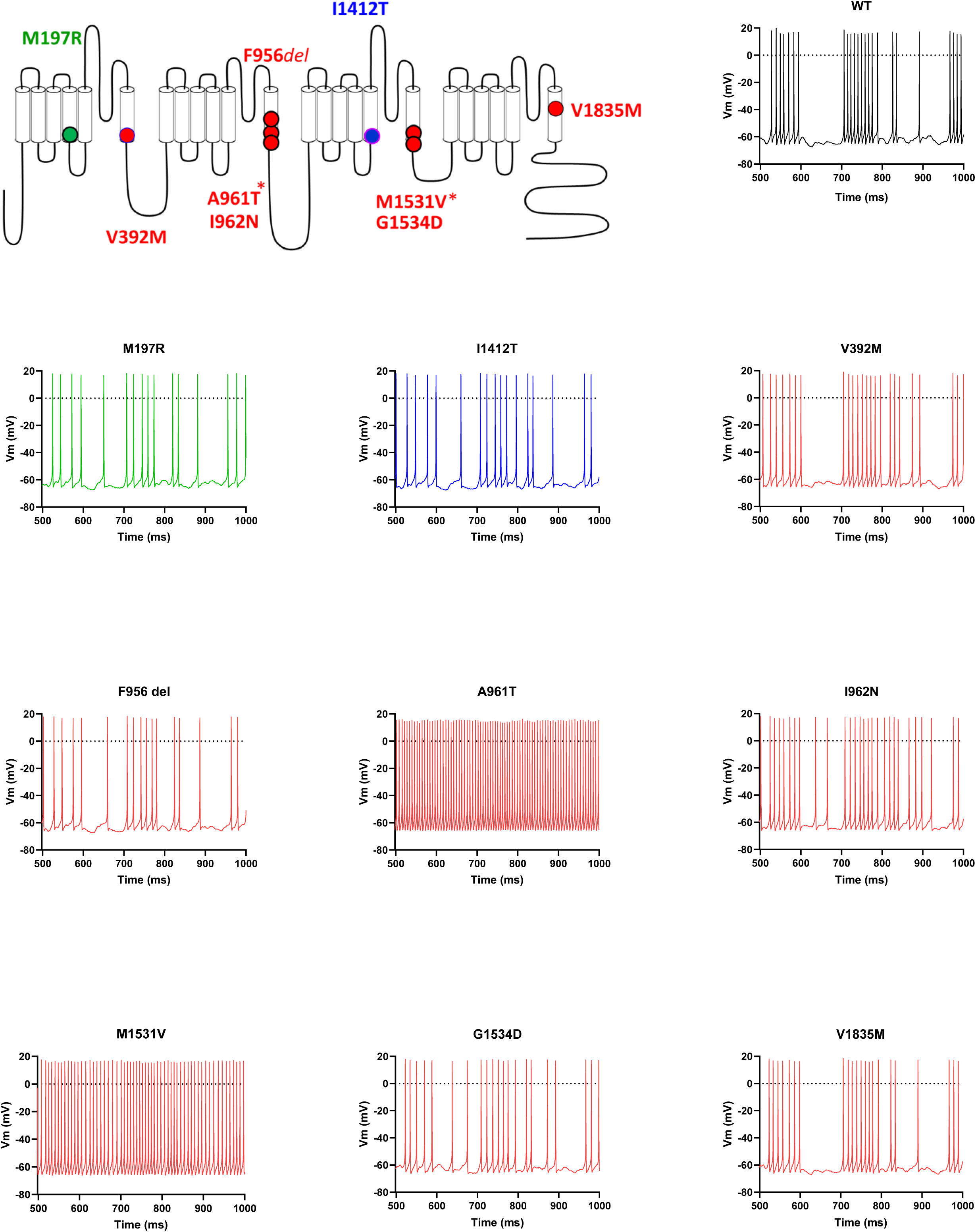
DCN modeling normalized for current density (MPC param.)

## Notes

### Competing Interest Statement

The authors have declared no competing interest.

